# Long-read cDNA Sequencing Enables a ‘Gene-Like’ Transcript Annotation of Arabidopsis Transposable Elements

**DOI:** 10.1101/2020.02.20.956714

**Authors:** Kaushik Panda, R. Keith Slotkin

**Affiliations:** Donald Danforth Plant Science Center, St. Louis, MO, USA; Division of Biological Sciences, University of Missouri, Columbia, MO, USA

## Abstract

High-quality transcript-based annotations of genes facilitates both genome-wide analyses and detailed single locus research. In contrast, transposable element (TE) annotations are rudimentary, consisting of only information on location and type of TE. When analyzing TEs, their repetitiveness and limited annotation prevents the ability to distinguish between potentially functional expressed elements and degraded copies. To improve genome-wide TE bioinformatics, we performed long-read Oxford Nanopore sequencing of cDNAs from Arabidopsis lines deficient in multiple layers of TE repression. We used these uniquely-mapping transcripts to identify the set of TEs able to generate mRNAs, and created a new transcript-based annotation of TEs that we have layered upon the existing high-quality community standard TAIR10 annotation. The improved annotation enables us to test specific standing hypotheses in the TE field. We demonstrate that inefficient TE splicing does not trigger small RNA production, and the cell more strongly targets DNA methylation to TEs that have the potential to make mRNAs. This work provides a transcript-based TE annotation for Arabidopsis, and serves as a blueprint to reduce the genomic complexity associated with repetitive TEs in any organism.

## Introduction

A consistent problem with the analysis of eukaryotic genomes is the complexity introduced by transposable elements (TEs). Thousands to millions of TEs are present in eukaryotic genomes, often nested in convoluted organizations. Analysis of these regions is cumbersome due to their repetitive nature (number of similar or even identical elements) and the fact that current TE annotations only describe the bare minimum of TE information. Even the best TE annotations consist of only three features: position on the chromosome, strand and TE type. In contrast, gene annotations are regularly based on transcript information, which is missing for TEs. These gene annotations describe the transcriptional start sites (TSSs), polyadenylation site, direction and splicing pattern, which provide genes with higher resolution in bioinformatic experiments compared to TEs. The lack of transcript information for TEs hampers downstream bioinformatics, leading many researchers to ignore these regions of the genome altogether.

A TE transcript-based annotation would enable research on TE expression: which elements are expressed, the transcript forms they generate, and when they are differentially expressed. However, two shortcomings have prohibited using RNA-seq information to generate a transcript-based annotation of TEs. First, short-reads generated by Illumina sequencing perfectly map to many TE locations, creating ambiguity regarding which TE is expressed. Second, TEs are subject to overlapping mechanisms that repress their expression, such as maintenance of epigenetic transcriptional silencing, small RNA-based chromatin modification and post-transcriptional silencing mediated by RNAi (reviewed in (Ozata et al., 2019; Deniz et al., 2019)). Without TE expression (due to TE silencing), TE transcripts cannot be captured for characterization.

We overcame both of these technical difficulties by producing a long-read transcriptome annotation in Arabidopsis plants that are deficient in multiple layers of TE repression. Previous research in Arabidopsis has identified key proteins that function in TE silencing. The DDM1 protein acts to condense heterochromatin and maintain the silencing of TEs, and subsequently the *ddm1* mutant results in a broad activation of TE expression (Hirochika et al., 2000)(Lyons and Zilberman, 2017). The Pol V protein complex acts in the RNA-directed DNA methylation (RdDM) pathway to reinforce DNA methylation at short TEs and re-silence active TEs (Lahmy et al., 2009)(Zhong et al., 2012)(Panda et al., 2016). The RDR6 protein generates double-stranded RNA, triggering mRNA degradation and TE small interfering RNA (siRNA) production via RNAi (Baeg et al., 2017; Nuthikattu et al., 2013). We combined mutations in these three genes to activate and stabilize TE transcripts, performed Oxford Nanopore Technology (ONT) full-length cDNA sequencing, and generated a first-of-its-kind genome-wide transcript annotation for TEs. Our annotation enables higher bioinformatic resolution of TE regions of the genome, allows focus on the potentially functional TE copies, and opens testing of long-standing hypotheses on the regulation of TE expression.

## Results

### Long-read transcriptome sequencing of TE-activated lines

We began by isolating total RNA, purifying polyadenylated mRNAs, and performing ONT sequencing of full-length cDNAs from five Arabidopsis genotypes (Figure 1A)(see Methods)(sequencing statistics in Supplemental Table 1). The genotypes selected included wild-type reference strain Columbia (wt Col), which has transcriptionally silent TEs, and the *ddm1* mutant with transcriptionally reactivated TEs (Zemach et al., 2013). We also combined the *ddm1* mutation with either *rdr6* or *pol V*, or both *rdr6* and *pol V* in the *ddm1/rdr6/pol V* triple mutant. We found that in wt Col very few reads (3.7%) overlap known TE annotations (Figure 1B), and the majority of these are genic transcripts that read-through a TE annotation (~84%). Removing the read-through of genic transcripts into TEs, we find only 2,719 reads are *bonafide* TE-initiated transcripts (0.59%, Figure 1C). Overall, only 0.26% of TE bases are covered by TE-initiated reads in wt Col (Figure 1D), confirming the efficient epigenetic suppression of TE expression in wt Col plants. The few expressed TEs in wt Col include AT2TE16945 and AT4TE03410 (*Sadhu* family non-LTR retrotransposons)(Figure 1E), AT3TE63065 (a *Copia* family LTR retrotransposon), AT5TE72200 (*TSCL* LTR retrotransposon) and AT5TE72580 (*AtSINE2A* non-autonomous element). Our data confirms previous findings that *Sadhu* family TEs and the *TSCL* element are expressed in wt Col (Rangwala et al., 2006; Chye et al., 1997).

**Figure 1.**
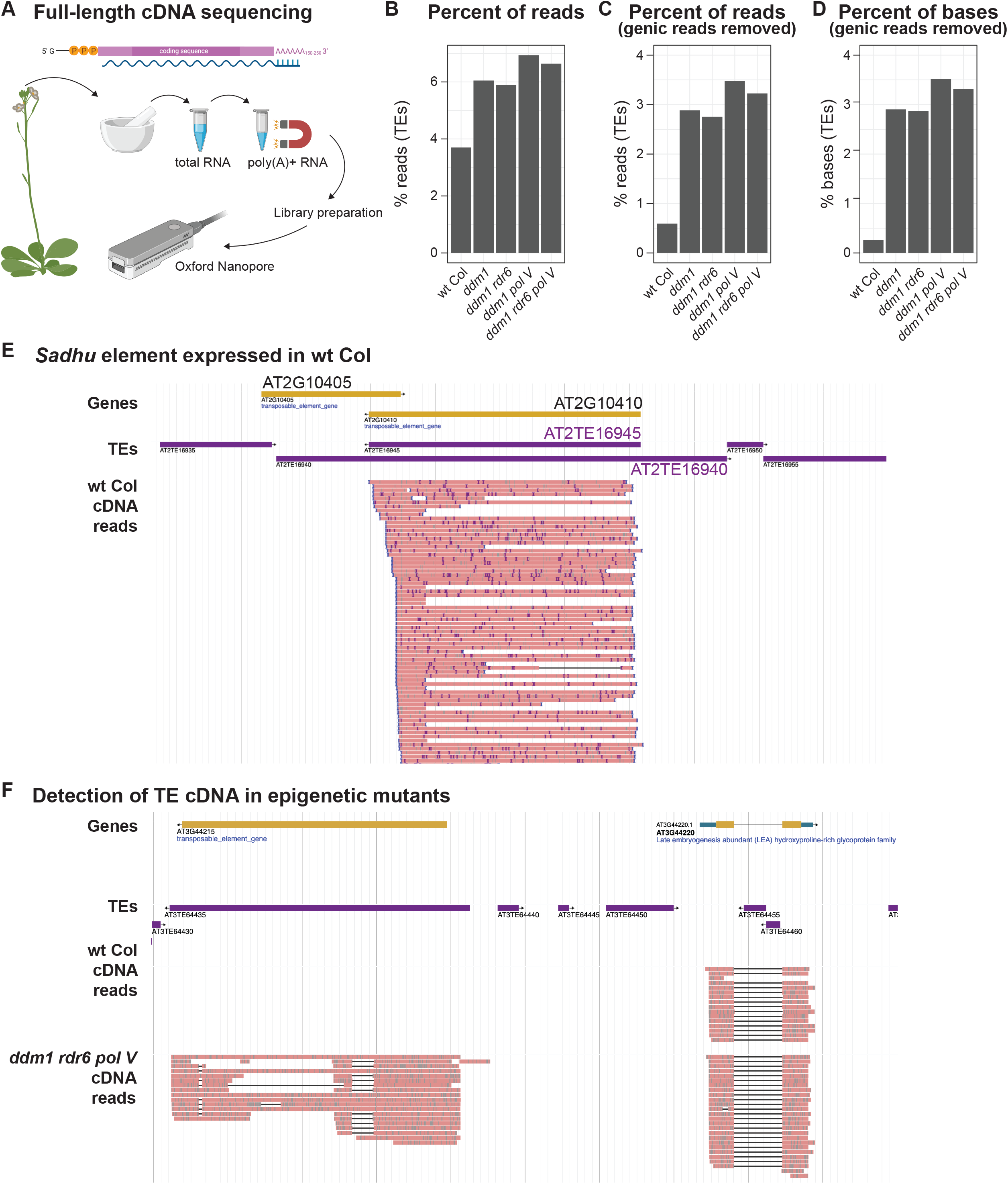
TE expression captured by long-read cDNA sequencing. **A**. Experimental workflow of ONT cDNA sequencing. See Methods for additional experimental details. Cartoon created with BioRender. **B.** Percent of reads that overlap a TE annotation in each genotype. **C**. Percent of reads that overlap a TE annotation, but with genic transcripts that read-through a TE removed to focus on TE-initiated transcripts. **D**. Percent of TE bases covered by reads, with genic transcripts that read-through TEs removed. **E**. Genome browser image of the expression of the *Sadhu* TE AT2TE16945 in wt Col. **F**. Genome browser image of the representative AT3TE64435 *AtCopia11* TE (left), which is only expressed in TE-activated mutants, compared to a gene expressed in both genotypes (right). In E and F genic exons are yellow, UTRs blue, TEs purple, and ONT cDNA reads are pink.

In the TE-activated mutants, we see a roughly 3-fold increase in the percent of TE reads and a 5-fold increase in percent of TE bases covered, and most of these are not read-through transcripts initiated by genes (Figure 1B-D). In the *ddm1* TE-activated mutant, we now detect transcripts originating from 31 *Gypsy* LTR retrotransposons, 24 *EnSpm* and 18 *Mutator* DNA transposons, and many others (Supplemental Table 2). As an example, transcripts from the *AtCopia11* element AT3TE64435 are only detected in TE-activated mutants (Figure 1F). Together, we demonstrate that epigenetic reactivation is required to capture TE transcripts for annotation.

### Transposable element transcript annotation

We performed additional sequencing on the *ddm1/rdr6/pol V* triple mutant to increase depth, as this genotype lacks three overlapping layers of TE suppression: transcriptional silencing via heterochromatin condensation / post-transcriptional mRNA degradation by RNAi / re-targeting of chromatin modifications via RdDM (Cuerda-Gil and Slotkin, 2016). We combined all 5,208,896 reads from all genotypes to generate a new transcript annotation for Arabidopsis, which was filtered specifically for TE-containing transcript models (see Methods). This provided 2,188 transcript models of 1,292 distinct TE annotations in the Arabidopsis genome, which include TE TSSs, transcript direction, splicing patterns and polyadenylation sites (Figure 2A). We did not perform *de novo* TE discovery, but rather layered whether a TE was expressed, transcript features and our TE annotation onto the existing high-quality TAIR10 TE list (Lamesch et al., 2011), which is broadly used by the community. Of the 31,189 TEs in the reference Arabidopsis genome, 24,431 (78.3%) showed no evidence of mRNA accumulation, 5,466 (17.5%) had at least one read but not enough to annotate a transcript, and 1,292 (4.1%) were expressed and transcripts were annotated. The annotated TEs are both euchromatic and pericentromeric, showing the same overall positional distribution as all TEs in the Arabidopsis genome (Figure 2B). To the community-standard TAIR10 TE list, we have added TE length, copy number, distance from the centromere, distance to the nearest gene, RdDM type (from (Panda et al., 2016)), whether or not the TE is expressed, TSS position, polyadenylation site, direction of transcription, transcript ID and transcript models (separate GFF file). These annotation files are Supplemental Data 1 and 2, and version controlled on GitHub (see Data Availability).

**Figure 2.**
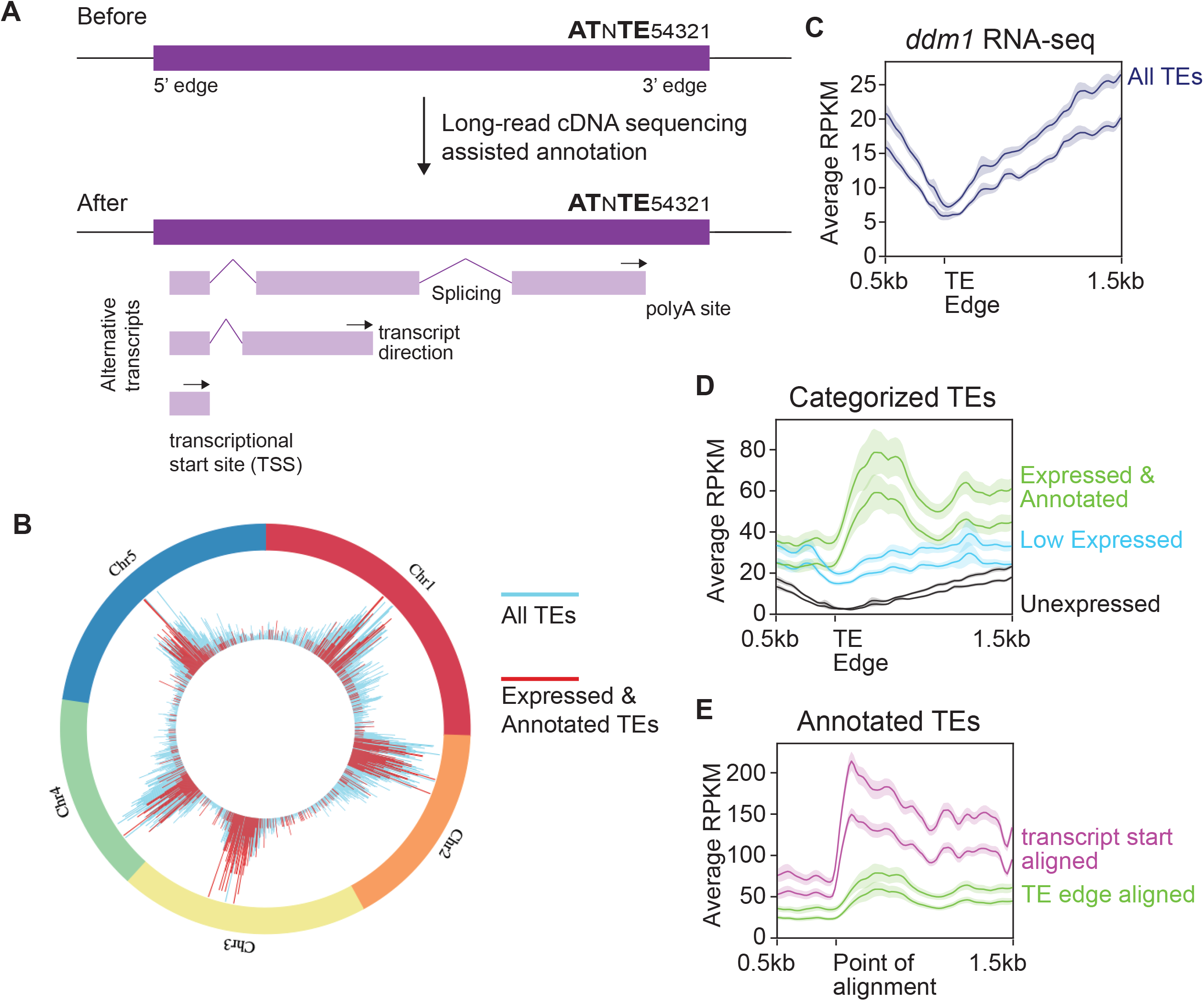
Genome-wide annotation of transposable element transcripts. **A**. Cartoon of TE annotated features before and after this work. **B**. Visualization of the chromosomal location of the TEs that are expressed and have been annotated in this work. **C**. Metaplot alignment of Illumina RNA-seq reads from *ddm1* to TE edges for all TEs. Y-axis represents the average reads per kilobase per million (RPKM). The two lines represent distinct biological replicates, the solid line indicates the moving average, and the translucent region is the standard error. **D**. The same as C, but the TEs are divided into expression categories based on our ONT cDNA sequencing. Low Expressed TEs did not produce enough reads for transcript annotation. **E**. The same as D, but the Expressed & Annotated TE class shown twice, once aligned by the TE edge (green line, same as D), or aligned by the now annotated TE TSS (purple line).

We next aimed to assay the quality of our TE transcript annotation. We used the quantitative nature of high-depth short-read Illumina RNA-seq from TE-activated *ddm1* plants (Oberlin et al., 2017) to measure the accuracy of regions we annotated as TE TSSs. Without our TE transcript annotation, TEs cannot be aligned by their TSS, and have therefore been previously aligned by their edges (Figure 2C). Expression of all TEs in *ddm1* aligned by their 5’ edge shows a reduction in transcripts at this point (Figure 2C), due to the large number of unexpressed short TEs. Dividing TEs into the expression categories defined by our transcript annotation shows that the “Expressed & Annotated’’ classification now displays a peak of expression close to the 5’ edge (Figure 2D). Importantly, our TE transcript annotation enables improved centering of TEs by their TSS rather than their edge. When the short-read RNA-seq data is mapped to the same set of TEs, either aligned by their 5’ edges or their now-annotated TSSs, we observe a sharper increase of expression and a higher density of mapped reads for the TEs aligned by their TSS (Figure 2E). Therefore, the independent Illumina RNA-seq datatype validates our annotation of TE TSSs.

### Improved TE annotations provide higher bioinformatic resolution

A major problem in TE bioinformatics is the frequent inability to differentiate distinct individual TE copies from closely related sub-family members (Shahid and Slotkin, 2020). The limited length of short-read sequencing generates multi-mapping reads that perfectly map to two or more locations in the genome. Several approaches have been taken to handle these multi-mapping short-reads, while the long-reads generated by ONT are unique to a single TE copy in the genome. To illustrate this point, we focused on the low-copy AtCopia93 *Evadé* sub-family of TEs (Mirouze et al., 2009). Our ONT data demonstrates the EVD5 element generates three transcript models, the two LTRs of EVD2 are transcribed, and EVD1, EVD3 and EVD4 are not expressed (Figure 3A). When we map short-read Illumina RNA-seq data, multi-mapping reads can be handled four different ways. First, we can use our TE transcript annotation to guide the reads, which accurately represents the transcripts that ONT detected (Figure 3A). Second, we can use only unique-mapping reads, but these report only the interior of EVD5 and miss EVD2 expression (Figure 3B). Third, the same result as Figure 3B is obtained if we use the uniquely mapping reads to guide the multi-mapping reads (Jin et al., 2015). Fourth, we can randomly distribute multi-mapping reads among the locations they perfectly match, but this falsely identifies EVD1 and the interior of EVD2 as expressed (Figure 3C). Of the most commonly used approaches to handle multi-mapping reads, unique and guided mapping underestimate the number of elements expressed, while the random mapping overestimates. This problem is exacerbated with higher copy TE families. Therefore, our TE transcript annotation helps guide multi-mapping reads to the correctly expressed individual elements.

**Figure 3.**
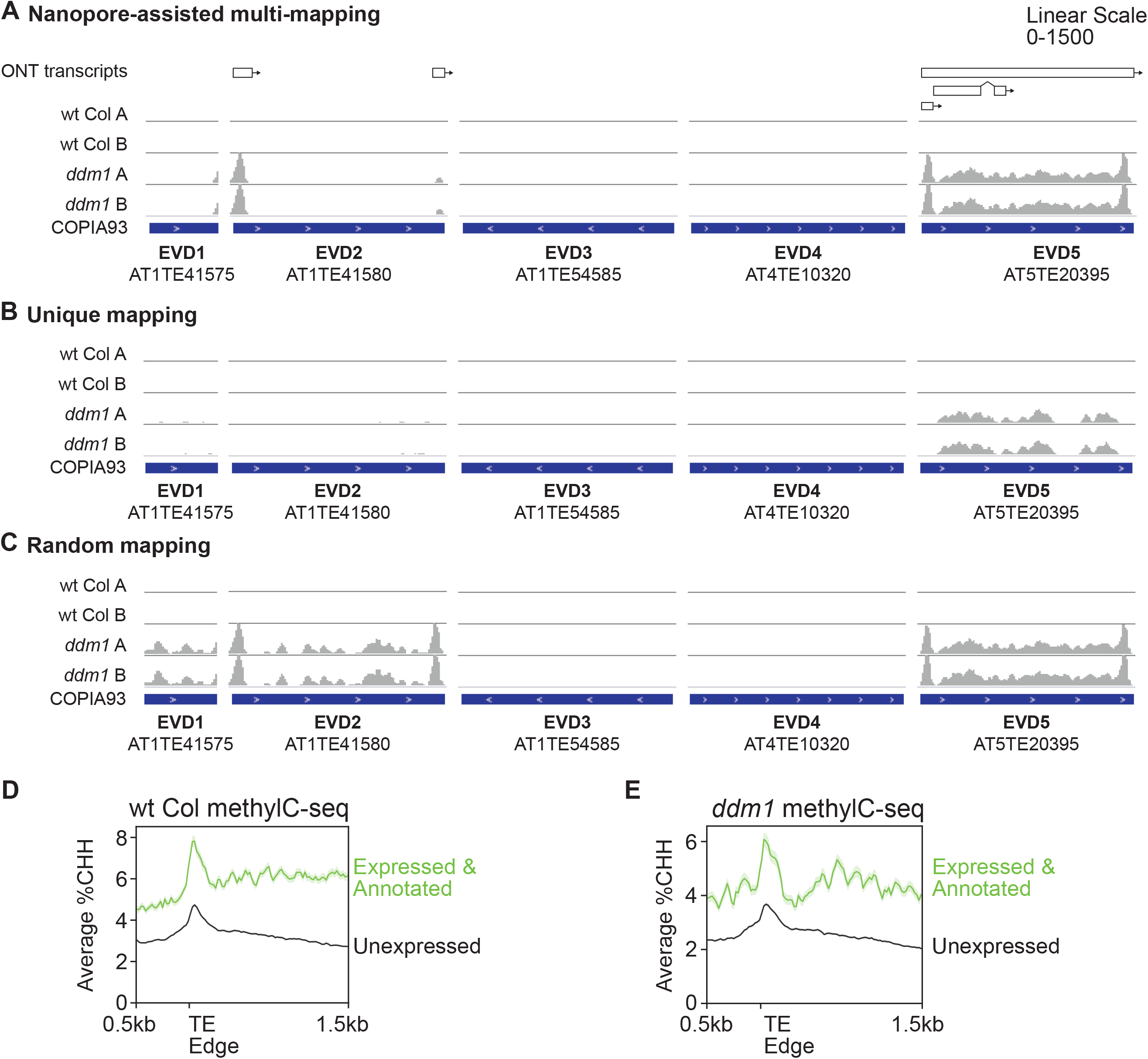
Increase in resolution of TE bioinformatics with improved TE annotation. **A**. ONT TE annotation of five *Evadé/AtCopia93* TE copies (top) and ONT-assisted mapping of wt Col and *ddm1* Illumina RNA-seq reads to the TE transcripts. **B**. Genome mapping of unique reads only. **C**. Genome mapping with multi-mapping reads distributed randomly. **D**. Genome-wide CHH context DNA methylation in wt Col assayed with Illumina methylC sequencing. TEs are aligned by their 5’ edge. **E**. Same as D, but with *ddm1* mutant plants.

In addition to mapping short-read RNA-seq, our TE transcript annotation provides resolution in other bioinformatic experiments such as genome-wide DNA methylation assayed by Illumina methylC-seq. Plant TEs have a known peak of CHH context DNA methylation (H=A,T,C) at their edges (Zemach et al., 2013), which is thought to reinforce the boundary between the TE and neighboring chromatin (Li et al., 2015). When TEs are transcriptionally silenced in wt Col plants, we show that our “Expressed & Annotated” TEs that have the potential to create mRNAs and our ‘Unexpressed’ TE category both have this peak of methylation at their edge (Figure 3D). This peak of CHH methylation remains in *ddm1* mutants (Figure 3E), demonstrating that the DDM1 gene is not responsible for defining the edge of TE chromatin boundaries. We previously hypothesized that TEs that create functional mRNAs would be targeted for higher levels of CHH methylation through the *expression-dependent RdDM* pathway (Panda et al., 2016). We found that the ‘Expressed & Annotated’ TE class defined by our transcript annotation has higher methylation at their edge in both wt Col and *ddm1* mutants (Figure 3D-E) compared to the ‘Unexpressed’ TE class. This finding supports our hypothesis and suggests that the cell can identify potentially mutagenic TEs and more strongly target them for repression. Thus, our transcript annotation provides a distinction between TEs based on their functional potential, which is important to test hypotheses on how TEs are targeted for repression.

### TE splicing is less accurate than gene splicing

Our TE transcript annotation allows the first transcriptome-wide investigation of TE splicing in plants. We compared splicing from TE-initiated transcripts with a set of genes that do not overlap TEs. There are 1,050 TE-annotated transcripts that do not have introns, while we focused on the 1,138 that have at least one intron and compared these to intron-containing genes. We found that the average number of introns is lower in TEs compared to genes (Figure 4A) and the TEs have on average larger exons (Figure 4B) and introns (Figure 4C). This suggests that TEs are subject to less overall splicing and potentially less splicing-based RNA quality control compared to genes.

**Figure 4.**
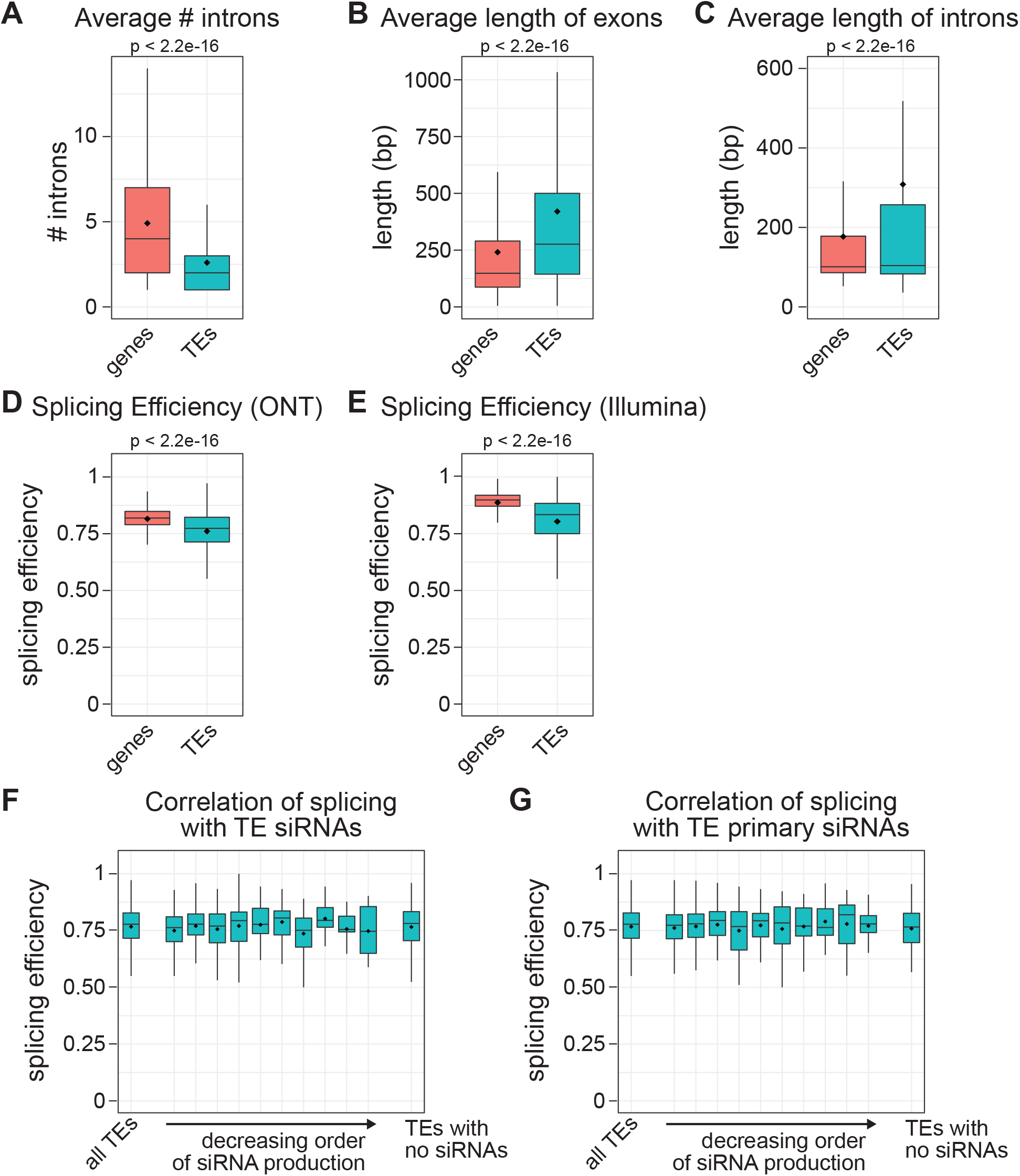
TEs have fewer introns and are spliced less efficiently than genes. **A**. Number of introns in genes and TEs with at least one intron. Box plots represent 25th and 75th quartile values with a line at the median, the mean is shown as a diamond and the whisker lengths represent the 10th-90th percentile. P-values are calculated by Welch’s two-sample t-tests. **B**. Length of exons in genes and TEs. **C**. Length of introns in genes and TEs. **D**. Accuracy of splicing defined by splicing efficiency based on ONT reads. **E**. Same as D, but using Illumina short-read RNA-seq reads to calculate splicing efficiency. **F**. Splicing efficiency of TEs ordered based on the amount of siRNAs generated in *ddm1/pol IV* double mutants. Splicing efficiency is based on ONT reads as in D, and ‘all TEs’ from D are shown as a reference. **G**. Same as F, but using only primary siRNAs generated in the *ddm1/rdr6/pol IV* triple mutant.

In two non-plant organisms, stalled spliceosomes and low efficiency of splicing have been shown to affect the quality control surveillance of TE mRNAs, specifically pushing TE transcripts into RNAi to generate primary small RNAs (Dumesic et al., 2013; Yu et al., 2019). The mechanism by which plant TE mRNAs are first degraded into primary siRNAs is currently enigmatic, so therefore we sought to test TE splicing efficiency in plants. We first used the raw ONT reads and compared these to our ONT-based transcript annotation to calculate the efficiency of splicing (see Methods), and found that TEs are not as efficiently spliced as genes (Figure 4D). Second, we compared Illumina short-read RNA-seq data to our transcript annotation and again found reduced splicing efficiency for TEs compared to genes (Figure 4E). We conclude that TE transcript splicing occurs less often and at a lower accuracy compared to genes.

To determine if the reduction in splicing efficiency found for TEs is correlated with the entry of TE mRNAs into RNAi, we examined siRNA production via small RNA sequencing. We first used a *ddm1/pol IV* mutant to assay siRNAs because of the abundant RNAi of some TE mRNAs that occurs in this mutant (Panda et al., 2016). We found no correlation between splicing efficiency and entry of specific TE mRNAs into RNAi (Figure 4F). Second, we performed the same analysis using small RNA sequencing data from a *ddm1/rdr6/pol IV* triple mutant, in which no secondary siRNAs are formed, allowing for the investigation of specifically TE primary siRNAs (Panda et al., 2016). We again found no correlation between TE mRNA splicing efficiency and the propensity of a TE mRNA to generate primary siRNAs (Figure 4G). This data demonstrates TE mRNA splicing efficiency does not trigger post-transcriptional degradation via RNAi, and overall proves the utility of a transcript-based TE annotation.

## Discussion

By performing long-read transcriptomics on a TE-activated line, we have captured full-length TE polyadenylated transcripts and used them to produce an improved TE annotation. We have demonstrated two biological consequences of this improved annotation. First, this method identifies the relatively small number (1,292) of individual TE copies in the genome capable of protein-coding expression, reducing the complexity of TE analysis. These transcripts can be used in future experiments to unambiguously map short-read RNA-seq data, identify the individual TE copies responsible for expression (similar to Figure 3A) and better calculate the differential expression of individual elements. Second, transcript-based TE information also enables current and future research testing hypotheses that require information on specific transcript features, such as TSSs or splicing patterns. We used this new TE transcript annotation to quickly test two standing hypotheses in TE biology. First, we found that TEs capable of expression and therefore potentially mutagenic activity are targeted more strongly by the cell for repression via DNA methylation. Second, we showed that inefficient TE silencing does not trigger RNAi on plant TE mRNAs. However, a limitation of our analysis is that it only tests the splicing efficiency on mRNAs, and not potentially inefficiently spliced RNA that is not polyadenylated. Nevertheless, with our added transcript-annotation based on the existing high-quality TAIR10 TE list, Arabidopsis now may have the best TE annotation of any multicellular organism.

In the current study, we used mutations as one way to activate TEs and produced a blueprint for how to improve TE annotations and handle complexity in the genome. In the future, different sources of TE activation can be used to improve existing TE annotations. Abiotic and biotic stresses are known to activate some TEs (reviewed in (Horváth et al., 2017)) and developmental relaxation of TE silencing (DRTS) occurs in some wild-type tissues such as pollen and endosperm (Anderson et al., 2019; Martinez and Slotkin, 2012; Slotkin et al., 2009). In Arabidopsis, we can pair these stresses and DRTS events with the mutant resources that we have taken advantage of to maximize the breadth of TEs covered. The stress and DRTS activation of TEs will be particularly useful for improving TE annotations in other plants that do not have available mutants, and where TE complexity poses a greater challenge.

## Methods

### Plants and materials

All plants used in this study were grown at 22°C in 16-hour light conditions in growth chambers. The specific alleles of all the mutants used are in Supplemental Table 1. Inflorescence tissue was collected and stored at −80°C before RNA isolation.

### Oxford Nanopore cDNA sequencing

Total RNA was extracted using TRIzol reagent and enriched for poly(A) mRNA using the NEBNext Poly(A) mRNA Magnetic Isolation Module. The cDNA-PCR Sequencing kit by Oxford Nanopore (SQK-PCS108) was used to prepare cDNA libraries from the poly(A) mRNAs. Briefly, 50 ng of poly(A) mRNA, as measured by Qubit RNA HS assay (ThermoFisher Scientific), was reverse transcribed using Superscript IV reverse transcriptase (Thermo Fisher Scientific). The cDNA was PCR amplified for 13-14 cycles with specific barcoded adapters from the Oxford Nanopore PCR Barcoding Kit (SQK-PBK004). The amplified cDNA was purified using AMPure XP beads (Beckman Coulter). Finally, the 1D sequencing adapter was ligated to the DNA before loading onto a SpotON flow cell (FLO-MIN 106D R9 version) in a MinION sequencer. MinKNOW 3.1.19 was used to run the sequencing.

### Read Processing

Raw fast5 reads were basecalled and demultiplexed using Guppy version 3.1.5+781ed57. Reads were mapped to the Arabidopsis Araport11 genome using Minimap2 (Li, 2017). Only primary alignments (~95%) were retained for analysis. A local instance of JBrowse v1.12.5 (Buels et al., 2016) was used to upload TAIR10 gene and TE annotation along with the nanopore primary alignments for browser shots shown in Figure 1E-F. Overlap of the reads with TAIR10 annotated TEs was used to calculate read percentage in Figure 1B. To facilitate identifying reads initiated within genes, Pychopper2 was used to orient reads. All the reads that initiated within annotated genic exons or 5’ UTRs and matching the annotated transcript direction were marked as ‘Gene’ reads and the remaining reads were overlapped with annotated TEs for calculation of TE-initiated reads (Figure 1C) and bases (Figure 1D).

### Transcript Annotation

The Pinfish pipeline by Oxford Nanopore was used to call *de novo* transcripts using long reads and default parameters (except a minimum cluster size of 5 was used). Similar to the analysis of individual reads, transcripts that originated within genic exons or 5’ UTRs and in the matching orientation were annotated as ‘Gene’ transcripts. Of the remaining transcripts, only those transcripts were annotated as TE transcripts if at least 25% of an exon overlapped with an annotated TE. The TE superfamilies represented by the transcripts for each genotype are noted in Supplemental Table 2.

For annotating TE transcripts, all reads from all genotypes were pooled including the higher depth sequencing of the triple mutant (*ddm1/rdr6/pol V*) before using the Pinfish Pipeline. The pipeline was run twice, first with a minimum cluster size of 3 and then again with 5. The reason we ran the pipeline additionally with a lower cluster size was to capture transcripts from TEs that did not have high expression. These two transcript files were merged and duplicate transcripts were removed. A total of 34,254 transcripts were identified.

### Transcript Assignment to Genes and TEs

First, the genes in the TAIR10 list annotated as TEs were removed to retain a pure ‘Gene-only’ list. Gene-initiated transcripts were removed as described above. Of the remaining transcripts, the ones that had at least one exon which overlapped at least 25% with an annotated TE was assigned as a ‘TE transcript’. A total of 2,188 transcripts representing 1,292 TEs were annotated as TE transcripts. There were many transcripts that overlapped with multiple TEs due to nested TE configurations. To assign transcripts to individual TEs the following criteria was followed preferentially: 1) If a transcript overlapped with only 1 TE, the transcript was assigned to that specific TE, 2) If there are more than 1 TE that overlap with the transcript, the TE(s) that overlaps with significantly longer section of the transcript (at least 3-fold greater than other overlapping TEs) was/were annotated to the specific transcript. 3) Finally, if more than 1 TE continues to be assigned to 1 transcript, then the TE with matching strand to the transcript direction was assigned the transcript. Following these criteria, a total of 1,936 transcripts were assigned to 1 TE each, and 252 transcripts were assigned to more than one TE.

### Updating the TAIR10 TE annotation

The columns added to TAIR10 TE list for this new version of TE annotation are not only based on transcript annotation (current study) but also on TE characterization (from (Panda et al., 2016): TE length, length category, subfamily copy number, copy number category, distance from the centromere, position category (euchromatic/pericentromeric), distance to the nearest gene, RdDM type when TEs are silent (wt Col) and when TEs are active (*ddm1*). For transcript annotation, four columns were added. First, the expression category of TE: “ExpressedAndAnnotated” if the transcripts are annotated as mentioned above; “LowExpressed” if no defined transcripts but at least 1 TE-initiating read was found in any of the genotypes; “.” If no TE-initiating read was detected in any of the genotypes. For the “ExpressedAndAnnotated” TE category, additional three columns were added: transcription start site, polyadenylation site and transcriptID. These columns may include comma separated values if more than one transcript is found for a specific TE. This TE annotation is Supplemental Data 1.

The transcriptIDs can be used to cross-reference with Supplemental Data 2 which is a GFF transcript annotation file that contains exon structure for each TE transcript. All TEs and “ExpressedAndAnnotated” TEs are used in the circular BioCircos plot (Cui et al., 2016) shown in Figure 2B.

### RNA-Seq validation

*ddm1* RNA-seq raw reads from GSE93584 were trimmed for adapters using Trimmomatic (Bolger et al., 2014), and mapped to the TAIR10 Arabidopsis genome using STAR (Dobin et al., 2013) with the following parameters: --runMode alignReads -- outMultimapperOrder Random --outSAMtype BAM SortedByCoordinate -- outFilterMultimapNmax 50 --outFilterMatchNmin 30 --alignIntronMax 10000 -- alignSJoverhangMin 3. DeepTools2 (Ramírez et al., 2016) was used to calculate normalized read count for each TE using a bin size of 20bp and generate the metaplots in Figure 2D-E.

### *Evadé* mapping strategies

Adapter trimmed RNA-seq reads for wt Col and *ddm1* (as mentioned above) were mapped to AtCOPIA93 (*Evadé*) copies using STAR with either unique strategy for multiple mapping reads (Figure 3B) or random strategy (Figure 3C). For nanopore-assisted multi-mapping, only the regions of AtCOPIA93 which produce transcripts were included as reference instead of all AtCOPIA93 copies (Figure 3A). The bam alignment file from all three mapping strategies was uploaded to IGV for the browser shots shown in Figure 3.

### Methylation metaplot analysis

Wt Col and *ddm1* methylC-seq processed files were used from GSE79746 to generate bigwig files using methylpy (Schultz et al., 2015). These bigwig files were used in deepTools to generate the methylation metaplots (Figure 3D-E) similar to the RNA-seq metaplots described above.

### Splicing Efficiency

To calculate splicing efficiency for nanopore cDNA reads, read depth of each position in the genome was calculated from the ‘all genotypes combined reads’ mapped file using samtools depth function (Li et al., 2009). Splicing efficiency was calculated as in (Yu et al., 2019) except that our calculation was based on the fall-off of read depth at exon/intron boundaries. For either TE or genic transcript files, 5’ splice site efficiency was defined as the ratio of depth of position upstream of the splice site and the added total depth of the positions upstream plus at the splice site. The 5’ and 3’ splicing efficiencies were averaged for all splicing events (multiple introns) in a transcript, and the transcript splicing efficiency was defined as the average of 5’ and 3’ splicing efficiencies. All individual transcript efficiencies were plotted in Figure 4D-E. For splicing efficiency of Illumina reads, depth calculated from *ddm1* RNA-seq reads from GSE93584 were used instead of nanopore cDNA reads.

### Correlation of splicing efficiency and small RNA production

To capture the TEs that produce siRNAs, we first identified TEs that produced at least 1 siRNA read per million in a *ddm1/pol IV* mutant. We used this mutant as it has the highest levels of TE RNAi (Panda et al., 2016). The TEs were sorted by the number of normalized siRNA production (averaged across two replicates) and divided into 10 categories. As a control, ‘all TEs’ and the TEs that do not produce at least 1 siRNA read per million were compared. Since RDR6 is involved in amplifying RNAi by generating sRNAs in a feedback loop, we aimed to combine *rdr6* mutation to remove secondary siRNAs and only investigate primary sRNAs. So we repeated the siRNA correlation analysis using siRNAs from *ddm1/pol IV/rdr6*. All siRNAs used were downloaded from GSE79780. Raw reads were trimmed for adapters, and mapped to Arabidopsis genome using ShortStack (Axtell, 2013)(fractional seeded strategy for multi-mapping reads). Samtools bedcov function was used to calculate normalized siRNA read count of each TE.

## Data Availability

All the raw data generated in this study are deposited in NCBI GEO database (GSEXXXXXX). The ‘Panda’ TE transcript annotation v1.0 files are Supplemental Document 1 and 2, and version controlled available at Github (https://github.com/KaushikPanda1/AthalianaTETranscripts).

## Acknowledgments

The authors thank Meredith Sigman for production of the *ddm1/rdr6/pol V* triple mutant line and Alex Harkness for technical assistance with ONT sequencing. This work is supported by NSF MCB-1608392 to R.K.S.

## Author Contributions

K.P. and R.K.S designed the research. K.P. performed the research and data analyses. K.P. and R.K.S. wrote the article.

---

**Supplemental Table 1.** Sequencing statistics and alleles used

|  | Flow cell 1 | Flow cell 2 |
| --- | --- | --- |
| Genotypes | wt Col, *ddm1*,<br>*ddm1/rdr6*,<br>*ddm1/pol V*,<br>*ddm1/rdr6/pol V* | *ddm1/rdr6/pol V* |
| Reads produced | 3,568,429 | 3,778,131 |
| Passed QC filter | 2,724,150 | 2,626,461 |
| Mean Read length (nt) | 841 | 805 |
| N50 | 972 | 948 |
| Mean Read Quality | 11 | 11 |
| Longest Read | 8,430 | 8,949 |

|  | wt Col | *ddm1* | *ddm1/rdr6* | *ddm1/pol V* | *ddm1/rdr6/pol V* flow cell 1 | *ddm1/rdr6/pol V* flow cell 2 |
| --- | --- | --- | --- | --- | --- | --- |
| Passed QC filter reads | 548,918 | 329,372 | 617,480 | 699,144 | 460,686 | 2,553,296 |
| Genome mapped reads | 505,792 | 292,004 | 560,153 | 636,879 | 400,927 | 2,208,924 |
| TE overlapping reads | 18,725 | 17,666 | 33,002 | 44,176 | 26,632 | 146,610 |
| TE initiated reads | 2,719 | 7,505 | 13,863 | 20,034 | 11,733 | 63,872 |

| Gene | Allele used |
| --- | --- |
| DDM1 | *ddm1-2* |
| RDR6 | *rdr6-15* |
| Pol V | SALK_017795 |
| Pol IV | SALK_128428 |

**Supplemental Table 2.**
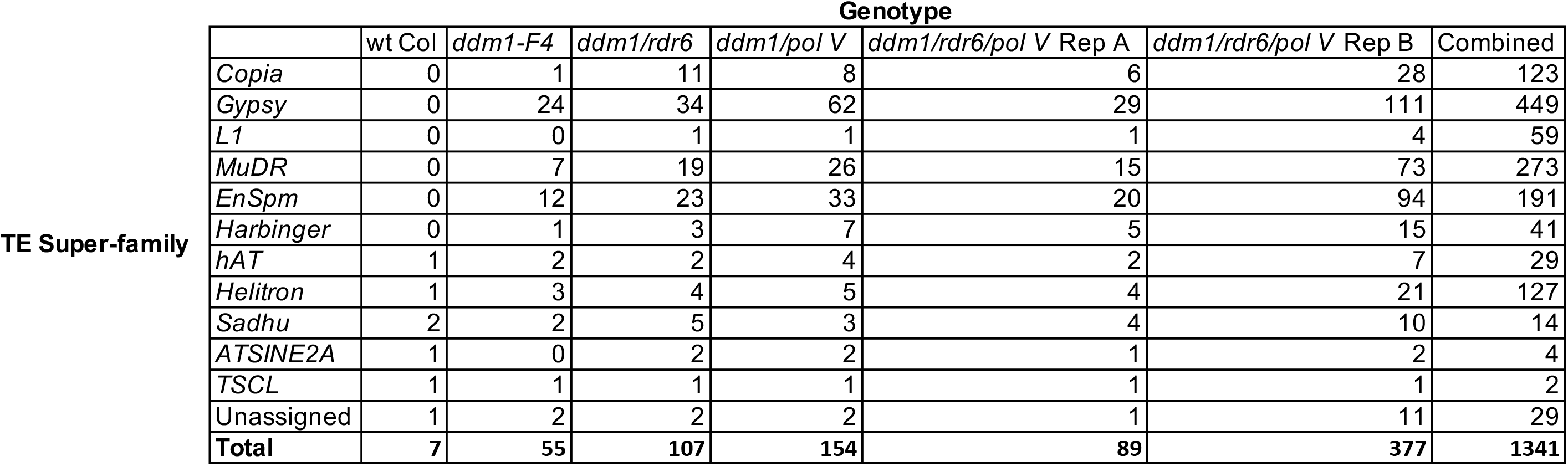
TE super-family expression detected in each genotype

|  | Genotype |  |  |  |  |  |  |
| --- | --- | --- | --- | --- | --- | --- | --- |
|  | wt Col | *ddm1-F4* | *ddm1/rdr6* | *ddm1/pol V* | *ddm1/rdr6/pol V* Rep A | *ddm1/rdr6/pol V* Rep B | Combined |
| *Copia* | 0 | 1 | 11 | 8 | 6 | 28 | 123 |
| *Gypsy* | 0 | 24 | 34 | 62 | 29 | 111 | 449 |
| *L1* | 0 | 0 | 1 | 1 | 1 | 4 | 59 |
| *MuDR* | 0 | 7 | 19 | 26 | 15 | 73 | 273 |
| *EnSpm* | 0 | 12 | 23 | 33 | 20 | 94 | 191 |
| *Harbinger* | 0 | 1 | 3 | 7 | 5 | 15 | 41 |
| *hAT* | 1 | 2 | 2 | 4 | 2 | 7 | 29 |
| *Helitron* | 1 | 3 | 4 | 5 | 4 | 21 | 127 |
| *Sadhu* | 2 | 2 | 5 | 3 | 4 | 10 | 14 |
| *ATSINE2A* | 1 | 0 | 2 | 2 | 1 | 2 | 4 |
| *TSCL* | 1 | 1 | 1 | 1 | 1 | 1 | 2 |
| Unassigned | 1 | 2 | 2 | 2 | 1 | 11 | 29 |
| **Total** | **7** | **55** | **107** | **154** | **89** | **377** | **1341** |

TE Super-family

